# Finding the signal in the Noise of Citizen Science Observations

**DOI:** 10.1101/326314

**Authors:** Steve Kelling, Alison Johnston, Daniel Fink, Viviana Ruiz-Gutierrez, Rick Bonney, Aletta Bonn, Miguel Fernandez, Wesley M. Hochachka, Romain Julliard, Roland Kraemer, Robert Guralnick

## Abstract

While many observations of species are being collected by citizen science projects worldwide, it can be challenging to identify projects collecting data that effectively monitor biodiversity. Over the past several years the allure of taking a “Big Data” approach has provided the opportunity to gather massive quantities of observations via the Internet, too often with insufficient information to describe how the observations were made. Information about species populations — where and when they occur and how many of them are there — (i.e., the signal) can be lost because insufficient information is gathered to account for the inherent biases in data collection (i.e., the noise). Here we suggest that citizen science projects that have succeeded in motivating large numbers of participants, must consider factors that influence the ecological process that affect species populations as well as the observation process that determines how observations are made. Those citizen science projects that collect sufficient contextual information describing the observation process can be used to generate increasingly accurate information about the distribution and abundance of organisms. We illustrate this using eBird as a case study, describing how this citizen science platform is able to collect vital contextual information on the observation process while maintaining a broad global constituency of participants. We highlight how eBird provides information with which to generate biodiversity indicators — specifically distribution, abundance, and habitat associations — across the entire annual cycle, even for populations of long distance migratory birds, a highly challenging taxon.

## Introduction

Monitoring biodiversity provides essential information with which to develop strategies for the conservation and sustainable use of resources. In most cases environmental monitoring programs rely on humans to collect field observations, because artificial intelligence systems are not yet able to classify organisms to species consistently (Kelling et al. 2013). Since governments and scientific agencies often lack resources to support long-term biodiversity assessment by professional scientists (Balmford and Gaston 1999, Bland et al. 2015), many organizations have developed citizen science projects that recruit the public to provide large quantities of monitoring data across large spatial and temporal extents (Amano et al. 2016, Danielsen et al. 2014, Pimm et al. 2014, Sullivan et al. 2014).

Thousands of projects engage the public in citizen science (see http://scistarter.com), with hundreds of these projects collecting observations of species (Theobald et al. 2015). Many projects take advantage of the global internet and the web to develop forms and mobile apps to gather data. Several projects achieve global participation, such as iNaturalist (http://www.inaturalist.org), which allows anyone to submit observations of any organism and manages a network of naturalists to identify the observations; REEF (http://www.REEF.org), which engages the diving community to monitor reef fishes; and eBird (http://www.ebird.org), which engages birders to gather bird observations. Combined, these citizen science projects are gathering millions of species observations annually (Chandler et al. 2016). eBird alone gathered more than 100 million records of individual species of birds (i.e., species observations) from 252 countries in 2017.

While citizen science has the potential to dramatically increase our biodiversity knowledge (Pimm et al. 2014), much care is required when interpreting citizen science observations for use in informing conservation and management actions. This is because citizen science can be motivated by varied objectives that may be at odds. Often a main objective is to inform science or conservation (Dickinson et al. 2010), which requires that a project be designed to collect accurate and usable data. However, frequently a second objective is outreach or education, i.e., a desire for a wide range of project participants to improve their scientific literacy and take an active role in conservation issues (Bonney et al. 2014, Jordan et al. 2012, Price and Lee 2013). There can be a tension between these two objectives as they could result in diverging strategies for improving data quality and increasing participant recruitment and motivation.

Biodiversity data collection methodologies occur along a gradient. At one end are structured surveys that emphasize strict protocols designed to meet specific objectives, and often require trained participants to sample at defined locations. At the other end of the gradient are unstructured projects that recruit participants from a wide range of expertise, have few requirements for data collection, and often do not collect information that can be used to control for data collection biases. Simple data collection protocols provide lower per-datum information content (Kery et al. 2010), but proponents contend that the sheer volume of data collected by a large number of citizen science projects is sufficient to meet scientific research objectives. However, if the data are gathered opportunistically and without any description of the sampling process, then even with large volumes of data, meaningful interpretations are limited in scope and the data are difficult to analyze and interpret (Conrad and Hilchey 2011, Kamp et al. 2016, Ottinger 2010). However, along this gradient there are projects that succeed in achieving broad public participation while gathering sufficient information to allow post-data collection analysis that controls for some known data collection biases.

The goal of this paper is to suggest best practices for improving the quality of species observation data gathered by citizen science projects. We emphasize the need for projects to gather data grounded within a scientific framework to produce accurate measures of the occurrence, abundance, and other attributes of species’ populations. We begin by reviewing key data-quality issues in citizen science projects that gather species observations. Next, we recommend several core fields of information that all citizen science projects should collect to address data quality issues. Our fundamental argument is that if citizen science projects collect a small set of basic information in addition to species observations, they can dramatically improve the scientific value of the information for the purpose of monitoring biodiversity.

### Data Quality Issues in Citizen Science Projects Focused on Species Observations

Regardless of where a project fits along the data collection methodology gradient, the method used to collect and record data must be considered and accommodated when project data are analyzed. The data are generated from a combination of two processes: 1) an ecological process that determines which species exist in a given location; 2) an observation process that determines which of the species that are present have been detected, identified, and reported. Structured surveys reduce the variation in the observation process by having rigorous protocols (Figure 1). Unstructured surveys can account for the observation process if they collect information sufficient to characterize and describe the process (Figure 1). When the two processes are confounded in the data, critical interpretations of ecological processes may be limited or misleading (Nichols et al. 2012). For example, an observed pattern in species occurrence may be related more to where sampling for that species occurred than to the actual occurrence of the species (Figure 2a). This is one of several known biases generated by unstructured surveys that need to be addressed, regardless of whether data are analyzed in a way that formally separates the ecological and observation processes (e.g., MacKenzie et al. 2006), or not (e.g., Fink et al. 2010).

**Figure 1.**
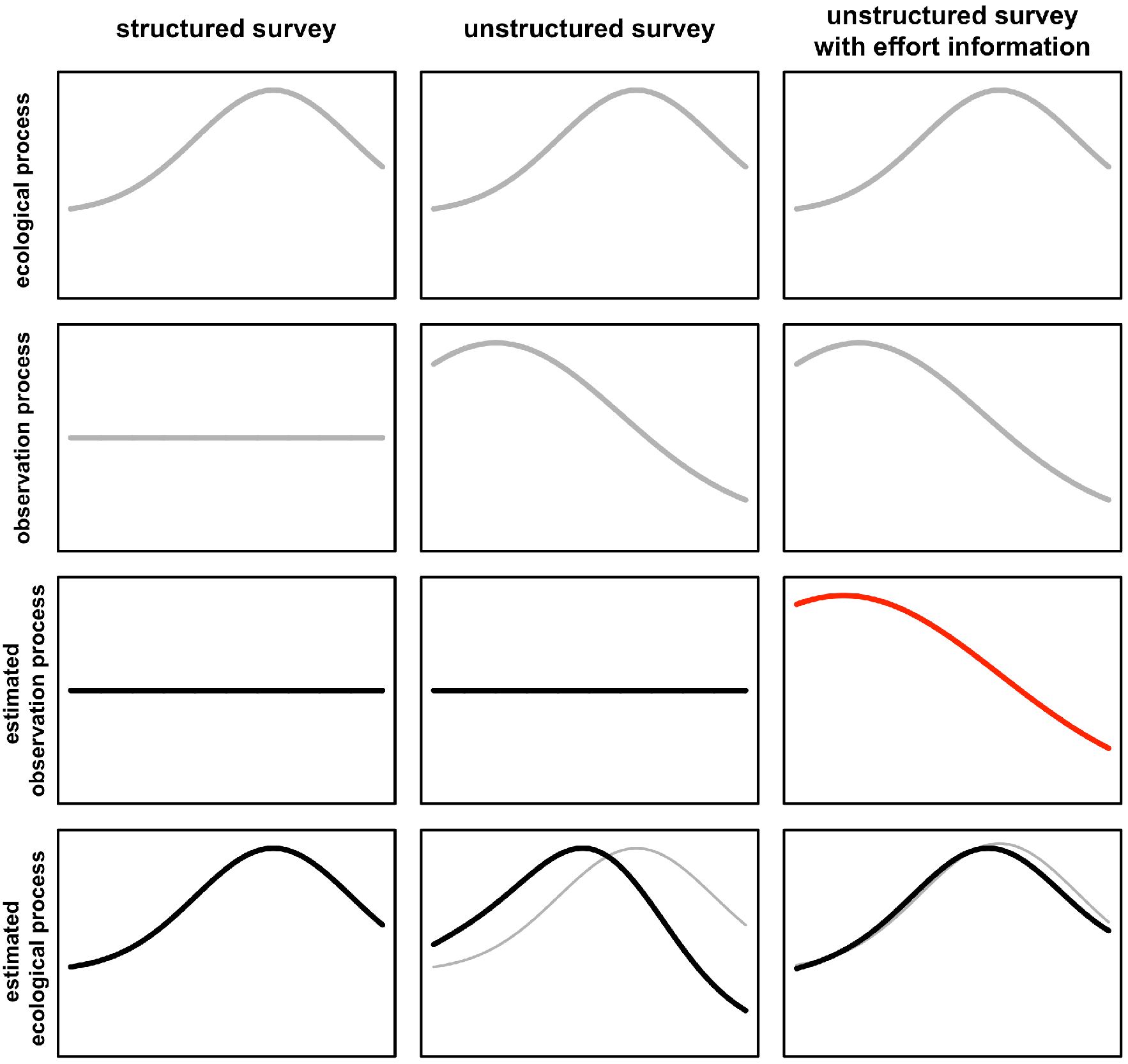
Schematic diagram of the ecological and observation processes. True processes are in grey, whilst estimated processes are in black and red. The ecological process (first row) shows the changing density of a species through a landscape (e.g., along a gradient from urban to rural environments). The second row shows the observation process. This is constant for a structured survey, which has consistent survey effort. The unstructured protocols have variable observations. In this example, consider more surveys nearer urban areas. The third row shows the estimated observation process. In unstructured surveys with no extra information (middle column) the observation process is assumed constant, so the observations are the product of the species density and survey density. In unstructured surveys with extra information (right column) it is possible to estimate the observation effort and this is indicated with the red line. The bottom row shows the density of species observations and the real ecological process (grey line). Both the structured survey and the unstructured surveys with effort information are able to recover something close to the true ecological process. The more closely the red line comes to the true observation process, the more accurately the ecological process will be uncovered. The unstructured survey without effort or bias information is not able to recover the true ecological process, because the ecological process is confounded with the observation process.

**Figure 2.**
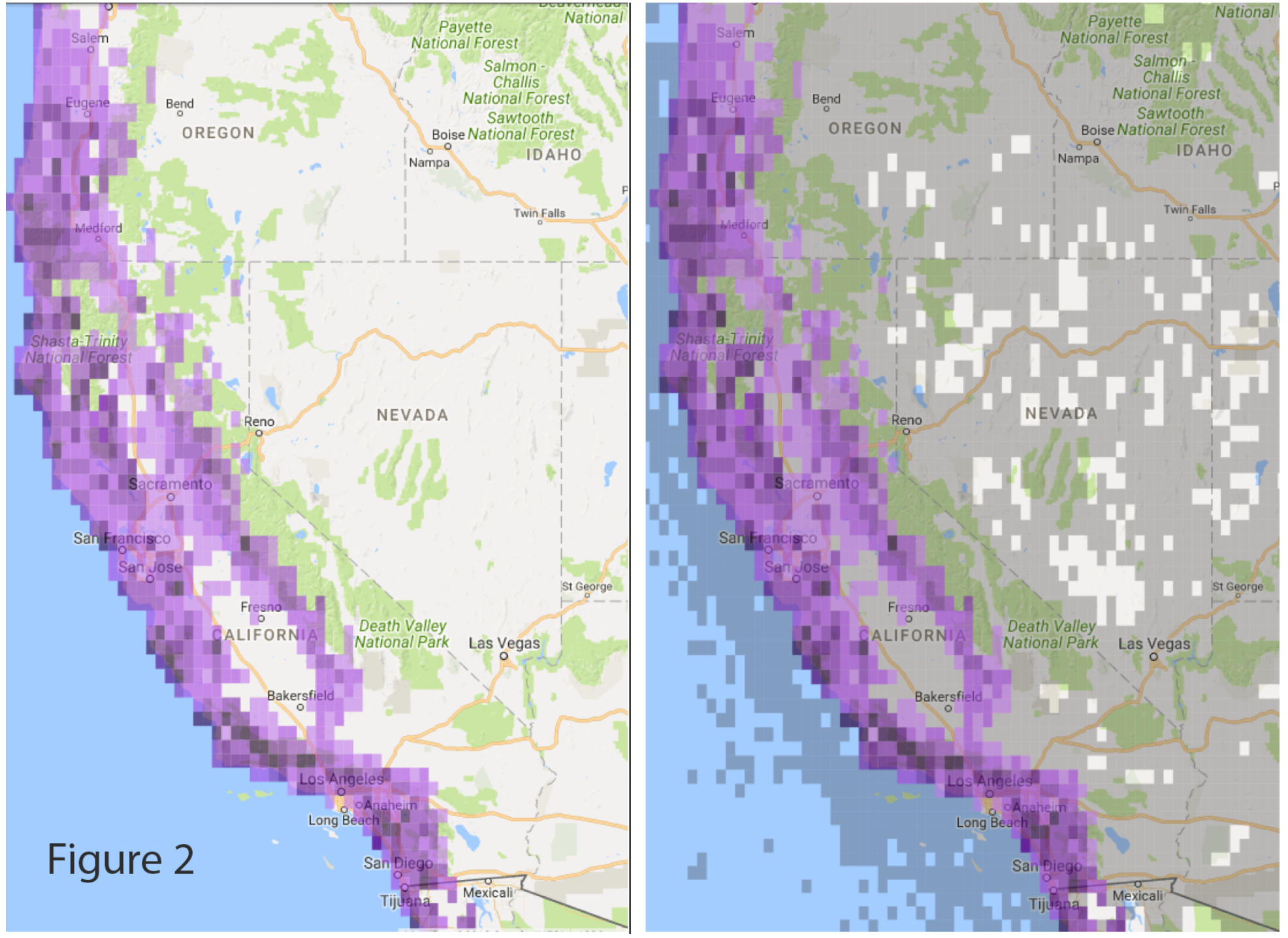
*left*. Presence-only locations where Wrentit (*Chamaea fasciata*) was observed in eBird. The darker the 20 km purple grid cells’ color, the higher the frequency of reports. Note in central California and Nevada Wrentits were not reported. Because these are presence-only data, it is unclear whether the map represents the true species range or sampling limitations. **Figure 2** *right*. Presence locations where Wrentit was observed in eBird (purple) and areas where observations were submitted but Wrentit was not observed (gray). This map provides much stronger evidence of species distribution.

The key step in accounting for variation in the observation process is to describe and control for known sources of bias. Biases can be separated into three major categories: 1) uneven sampling effort over space and time (Figure 3) (e.g., (Geldmann et al. 2016); 2) uneven detectability and identification of organisms (Figure 4) (e.g.,(Kery and Schmid 2004); and 3) uneven observation skill across participants (Figure 5) (e.g., (Crall et al. 2011, Delaney et al. 2008). Structured projects with rigorous data-collection techniques address these biases during data collection or through protocols that facilitate bias removal during analysis. Many unstructured projects, however, do not deal with these biases at all. For example, the majority of citizen science projects document the ecological process, but do not document observation process. We suggest that if unstructured citizen science projects collect sufficient information about the observation process, then controlling for these observation biases can be addressed during data analysis (Bird et al. 2014, Fourcade et al. 2014, Stolar and Nielsen 2015). We now review the differences between structured and unstructured projects.

**Figure 3.**
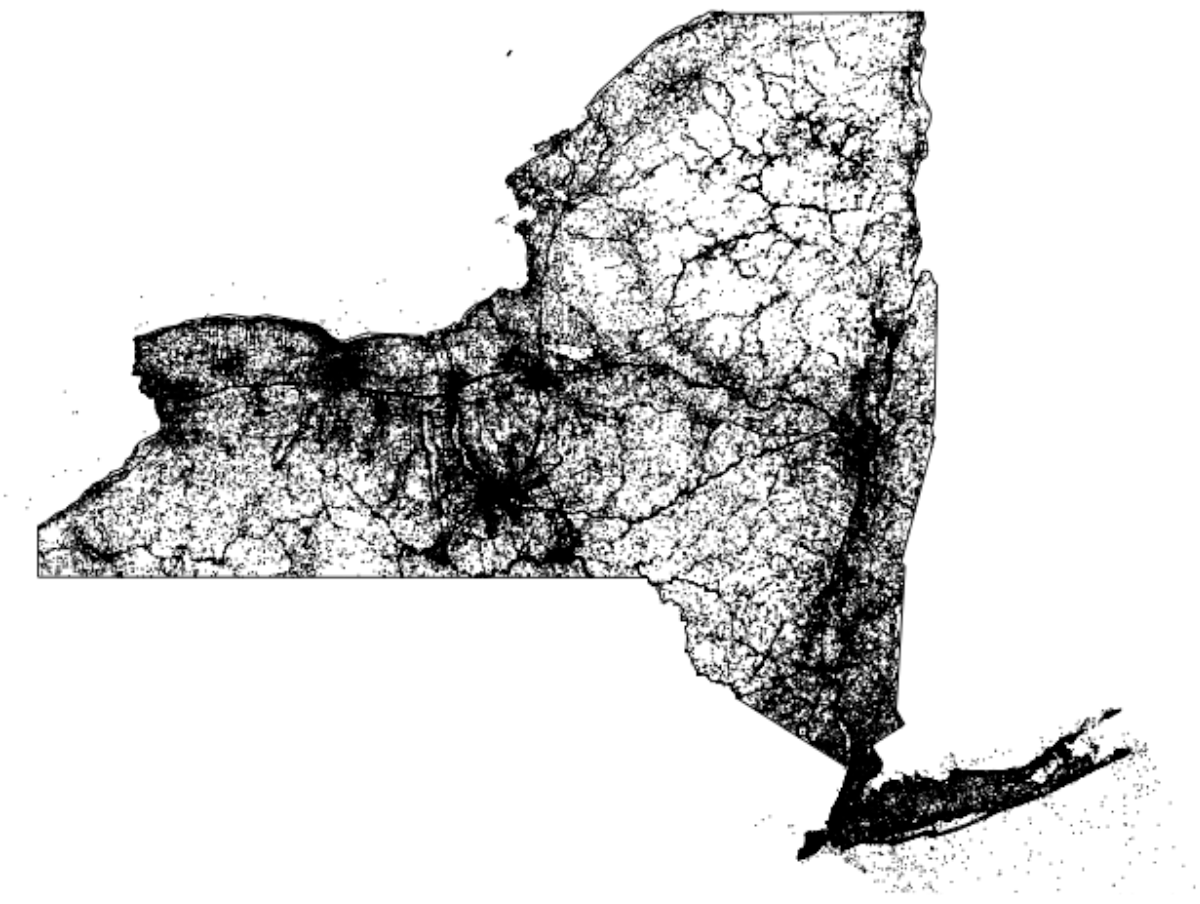
Locations in New York State where eBird participants have submitted observations. There are 214,000 unique locations where observations have been submitted. Dark areas are regions with high densities of observations. Note how dark areas are often correlated to areas with high human population densities.

**Figure 4.**
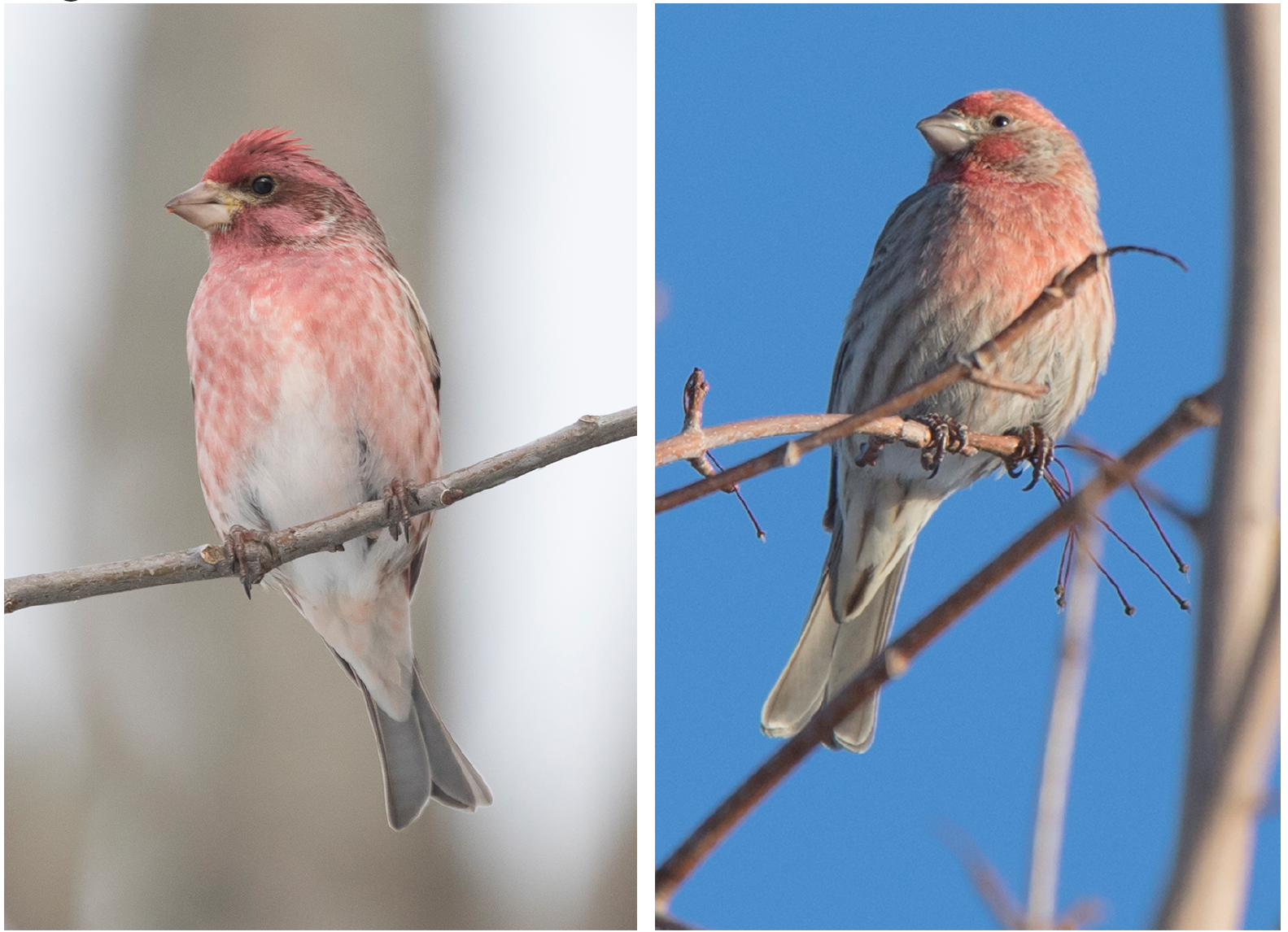
Identification of species is a difficult and confusing task. One of the biggest mistakes made by observers in North America is confusing male Purple Finch, *Haemorhous purpurus*, (left) and House Finch, *Haemorhous mexicanus*, (right).

**Figure 5.**
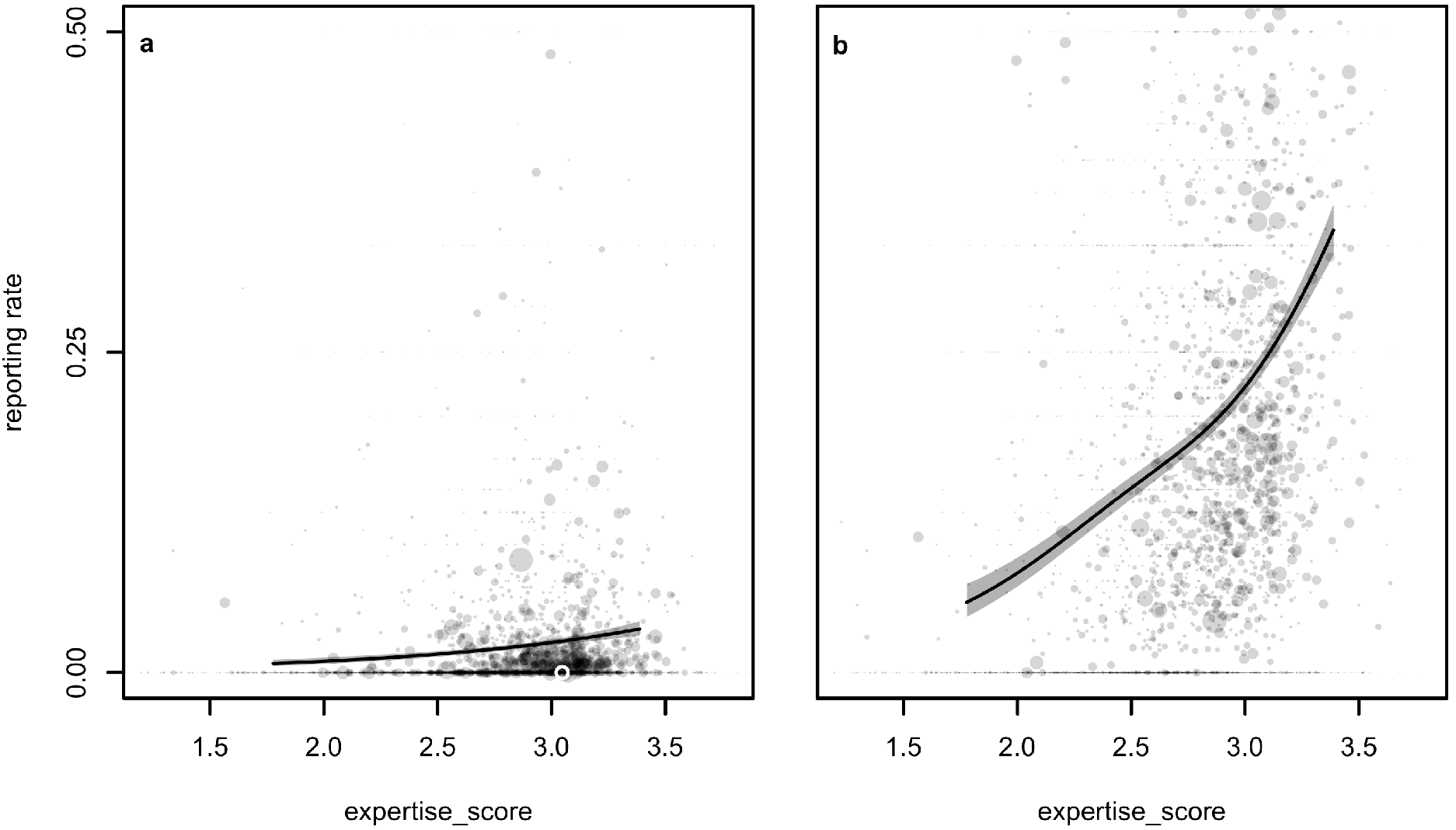
Reporting rate of a) Purple finch and b) House finch on checklists increases with observer expertise. Reporting rate is higher for house finch but increases with expertise for both species. Lines and grey boxes show fitted relationships and 95% confidence intervals. Each dot is the expertise and reporting rate of an individual observer, scaled by the number of checklists they submitted in the region and time period.

*Sampling Effort.* Structured data collection protocols preselect locations and clearly describe how and when data are to be collected. Sampling typically occurs during a specified time of the year, and locations are often selected using stratified sampling techniques (Albert et al. 2010). Data collection effort is tightly controlled by protocols that determine when, for what amount of time, and in what ways an observer collects observations. These approaches minimize bias and often standardize effort and sampling intensity. Examples include numerous bird atlases such as the Breeding and Wintering Birds of Britain and Ireland (Balmer et al. 2013) and the Swiss Biodiversity Monitoring Program (BDM_Coordination_Office 2014).

Unstructured data collection protocols allow flexibility in where an individual participates and do not have specific requirements about how and when data are to be collected. Data can often be collected any time of year and day. There are usually no data collection protocols and observers are required to report only species they observed. These unstructured projects have had only marginal success in using their data to monitor biodiversity, without significant assumptions being made. An example of an assumption is post-processing species presence data into a comprehensive species “list,” which has the potential to be a useful surrogate for effort expended in biodiversity surveys (van Strien et al. 2010). There is a class of unstructured projects that do collect information on the observation process. These projects have few protocol requirements, but they do collect information to identify the date, location and observer, and encourage the reporting of the start time and duration they collected observations, and the distance they walked collecting observations. If sufficient information on the observation process is recorded, then biases in the observation process can be addressed during analysis.

*Uneven detectability of organisms.* All biodiversity data collection projects suffer from the challenge of uneven species detectability (Johnston et al. 2014). This issue involves accounting for both false positives — misidentifications of observed organisms, and false negatives—failures to report species that were present. Some false positives can be filtered out as anomalies that fall outside the norm of occurrence for a species at a particular time or space. However, false positives can also be misidentifications of expected species. An example of this challenge is identification of Purple Finch and House Finch (Figure 4), which are very similar in appearance. False-positive rates are difficult to identify and correct for in statistical analyses without additional sources of information, such as testing participants about species identification abilities or using validation approaches that estimate observer skill levels (Kelling et al 2015b, Ruiz-Gutierrez et al 2016). In contrast, false negative errors can be corrected for, typically through the direct estimation of detection probabilities (i.e., the probability of detecting a species when present (MacKenzie et al 2003).

Structured surveys that employ trained participants using explicit data collection protocols can often directly control for species’ detectability. For example, many bird surveys require participants to make repeat observations (e.g., replicates used to estimate detection probability) or to collect observations at times of the year or times of the day when organisms are most active to maximize the opportunity for an observer to detect an organism.

Unstructured surveys must collect more than basic encounter information if they wish to measure variation in detection rates of species. Unstructured projects that require observers to indicate when they have contributed a complete list of all the species they detected allows an analyst to assume that any species not reported is a non-detection for that sampling period (Figure 2b). Complete information on the detection and non-detection of a species can be combined with sampling effort information to allow for the analytical control of detection rates and inference of variation in the probabilities of true absence (Fink et al. 2010).

*Uneven observation skills across participants.* While various modeling techniques can account for variation in detection rates of a species, they cannot distinguish differences in detection rates among observers without gathering and maintaining information about those observers. Structured surveys often use participants who are paid technicians or experienced volunteers. Unstructured projects typically have few barriers to participation resulting in more substantial variability in participant skills in detecting and identifying species. Regardless of whether a project engages skilled participants or is open to anyone, a key feature of data collection and management should be the association of participant identifiers with the observations that each participant collected, which allow analysts to monitor the relative skills of participants and their activity. For example, eBird indexes observer variability through species accumulation curves, which describe how the total number of species reported increases with increasing time spent in collecting observations and how this rate of increase varies among observers. These differences in species accumulation curves vary among observers and provide a measure of observer skill (Kelling et al. 2015a). They can be used as a post-hoc data-derived measurement of observer skill levels that improves ecological inference (Johnston et al. 2018).

### Improving the quality of species occurrence data gathered by Citizen Scientists

To control for the known biases outlined in the previous section, we argue that citizen science projects should be rooted within a survey design framework that includes several basic data collection principles that provide a solid foundation for data analysis. We build upon existing recommendations for biological monitoring programs that collect sufficient information on the sampling event that can be used within a statistical analyses framework (Yoccoz et al. 2001). The goal is for anyone designing a citizen science project to not only determine what is to be observed, but to describe the context in which each observation will be made. The answers to the following questions define the associated data-collection protocols needed for data analysis:

1. <u>Why</u> is the project being conducted? Every citizen science project should have a clearly articulated purpose based on either a research question or specific monitoring agenda to provide guidelines on what should be observed and how data need to be collected (Bonney et al. 2009). Any disconnect between the goals of the survey and how the data are collected will limit project success. For example, if the goal of a project is to relate species occurrences with land cover variables, the spatial resolution of data collected must be clearly defined. Imprecise information regarding where observations were made will lead to faulty land cover and observation relationships. We recommend that the goals of a citizen science project be clearly defined, even if relatively broad, so that the proper data collection methodology can be developed.
2. <u>What</u> is observed? Most biodiversity monitoring projects identify a specific taxonomic scope (which taxa are targeted), because developing protocols that effectively gather observations of one taxon (i.e., mammals) would be different from another (i.e., stream invertebrates). Often projects require an observer to report a checklist of all the species they identified within a taxon, sometimes including counts. Recording all detected species (within a predefined list of species/taxa) creates what is often known as a “complete list,” which provides information on species detected and species not detected. Non-detections can be used to estimate detectability and to infer species absences. We recommend that the taxon scope be clearly defined to ensure that the particular framework on how observations are collected can be developed. We also recommend the use of “complete lists” (even if on a subset of the full list of species within a taxon) so that non-detections can be used in subsequent modelling (Guillera-Arroita et al. 2015).
3. <u>Where</u> are the observations collected? Environmental factors not only constrain where species can live, but the habitat or weather conditions can affect the detectability of organisms. The detectability can be altered because habitat and or weather make it more difficult for observers to see, hear, or identify species; or because habitat and weather influence behavior of animals. Structured biodiversity monitoring programs gather species occurrence information only at preselected locations. However, many citizen science projects allow the observer to select the location rather than enforcing pre-selected locations. The advantage of user-selected areas is that recruitment of participants is easier. However, observers typically want to go to natural areas with high biodiversity (Tulloch et al. 2012) or to locations near where they live (Figure 4), which can result in spatial imbalance of observation locations. This problem can be partially mitigated via increased rates of participation and adjusting weights of observations made from high-density locations. Regardless of how a location is selected, knowledge of precise locations of observations allows for connections to be made with data on environmental conditions, which are now often assembled from remote-sensed datasets. We recommend that the locations where data collection events occur are identified at high spatial resolutions, and that the projects recommend locations where increased observer effort would benefit the project.
4. <u>When</u> are the observations collected? Species detectability can change within a day or seasonally as behavior of animals change or plants alter their appearance. For example, daily flight patterns of butterflies are determined by local weather conditions. Many biodiversity monitoring programs collect observations only at a particular time of day, or during specific days or months of a year. Although some unstructured citizen science projects also have adopted this approach, others permit year-round participation. There are advantages to both approaches. Limiting timing of collection can be problematic, especially when the timing of events such as migration or nesting change from year to year. In addition, limiting sampling events to a short time frame in a given year may limit the participation of volunteers. Recording of the sampling process (e.g., date, time, and duration of observations) allows for a subset of data to be extracted for specific times of year of interest and for control of the amount of effort made during a data collection event. We recommend that the timing of data collection be matched to the goals of the project, and that the date, start time, and duration of the sampling process is recorded.
5. <u>Who</u> is making the observations? Structured monitoring programs often rely on trained technicians for data collection, or have dedicated, long-standing participants, while most unstructured citizen science programs do not restrict participation and have a range of participants from those who are very dedicated to others who submit data only occasionally. While participants can develop enormous expertise in gathering information through participation in citizen science projects (Kelling et al. 2015a), volunteers as well as trained technicians can exhibit tremendous variation in their detection and classification skills. Regardless of the monitoring project, it is important to know who collected each observation. In this way bias related to variability in observer expertise can be estimated. We recommend that the data management framework retains information about who collected the observations, by using a code that is unique to an individual observer.
6. <u>How</u> are the observations collected? It is critical to report information about the collection process needed for documenting effort and completeness of a survey — time spent observing, estimates of area surveyed, and any other variables that have a strong effect on the detectability of species in the survey (Kery et al. 2010). A high level of data quality can be maintained in the absence of a specific sampling design and protocol in unstructured projects if the same information fields are reported clearly during each observation event made by a volunteer. When a project is gathering species observations, volunteers should accurately provide the location where observations were made and any distance traveled when making observations; the date; the time; and the duration of time over which they made observations. This information can be used to filter the data to produce a high-quality dataset that is more similar to data collected by structured monitoring programs. For example, data can be filtered to only those records that resemble a standard “protocol,” such as single-point observations made between 5-10 minutes for a given region and time of year. We recommend that sufficient information is gathered to clearly record the observation process regardless of how flexible they observation process might be.

### Biodiversity analysis with unstructured data from eBird

<u>*Overview:*</u> eBird engages more than 400,000 volunteers to report bird observations based on how bird watchers typically observe birds, i.e., units of data collection are “checklists” of zero or more species including a count of individuals of each observed species (Kelling et al. 2015b). As of April 2018, 28 million “complete checklists” containing 461 million observations of bird species had been submitted to eBird since the project began in 2002.

In order for the eBird observation process to be clearly articulated, the six questions that describe how observations are made must be clearly described.

1. <u>Why</u> is eBird being conducted? eBird collects data that will be used to estimate the distribution, abundance, and trends of bird populations by taking advantage of the global network of bird enthusiasts who submit their observations to a central data repository (Sullivan et al. 2014).
2. <u>What</u> is observed? eBird gathers a complete list of all bird species observed along with counts of individuals of each species during a data collection event.
3. <u>Where</u> are the observations collected? eBird participants can select where they make their observations. All locations are georeferenced either through mapping tools provided on the eBird website, or, more accurately, through the GPS system available on mobile phones and used by the freely available eBird App.
4. <u>When</u> are the observations collected? eBird allows observers to record observations at any time of day or year.
5. <u>Who</u> is making the observations? Anyone can participate in eBird. However, individuals must register and login to eBird whenever they submit a list of birds. The eBird database links all observations to the registered individual.
6. <u>How</u> are the observations collected? The location, date, and start time are recorded for all collection events. Prior to submission of species observations, participants must also record whether they are submitting a list of all the birds they detected and identified, which allows analysts to infer the absence of species not reported. Additionally, participants are encouraged to record search effort information - the duration and the distance traveled during the data collection event. Together, all of this information comprises a “complete checklist,” which is the foundation for much of the analysis of eBird data.

Translating eBird checklists into useful biodiversity information describing patterns of species occurrence and abundance in space and time are done through Species Distribution or Niche models (SDMs). These statistical models estimate the distribution or abundance of a species by estimating relationships between the observed patterns of species occurrence and data describing the processes that give rise to these observations (Franklin 2009). The data describing this process includes environmental data, often collected via remote sensing, which describe important ecological processes, e.g., habitat selection, which determine the environments most likely to be occupied by a species, and data that describe important aspects of the observational process, like the information contained in eBird complete checklists. When the goal is ecological inference, the effects of the observation process are sources of bias. If these sources of bias cannot be controlled during data collection, then data describing these biases are essential to account for them during analysis (Figure 1). SDMs provide the analytical framework where ecological signal can be separated from observational noise.

Data from eBird complete checklists in combination with remote sensing data are used to create a series of biodiversity indicator data products for a species. These include range-wide, seasonal relative abundance estimates (Figure 6), weekly relative abundance estimates (Figure 7), and weekly estimates of the relative importance of land and water cover classes to species occurrence (Figure 8). The range-wide, seasonal relative abundance estimates are meant to show the population of each species across its entire distribution. The inclusion of relative abundance identifies the core range of the species. The weekly relative abundance estimates show where the species occurs and its relative abundance for every week of the year. The estimates also show regions where the species does not occur and locations where eBird does not have sufficient data. The habitat plots show the seasonally and geographically varying habitat associations across the entire life cycle of a species. These indicators can be used to contrast regions, seasons, and species, or to compare the expected costs of management decisions. For example, modeling how bird populations change throughout the year has uncovered seasonally complex species-environment relationships (Zuckerberg et al. 2016), identified novel aspects of habitat associations that can impact bird populations during migration (La Sorte et al. 2017), and identified seasonal resources needed for supporting bird populations during critical stages of their life history (Johnston et al. 2015, Reynolds et al. 2017).

**Figure 6.**
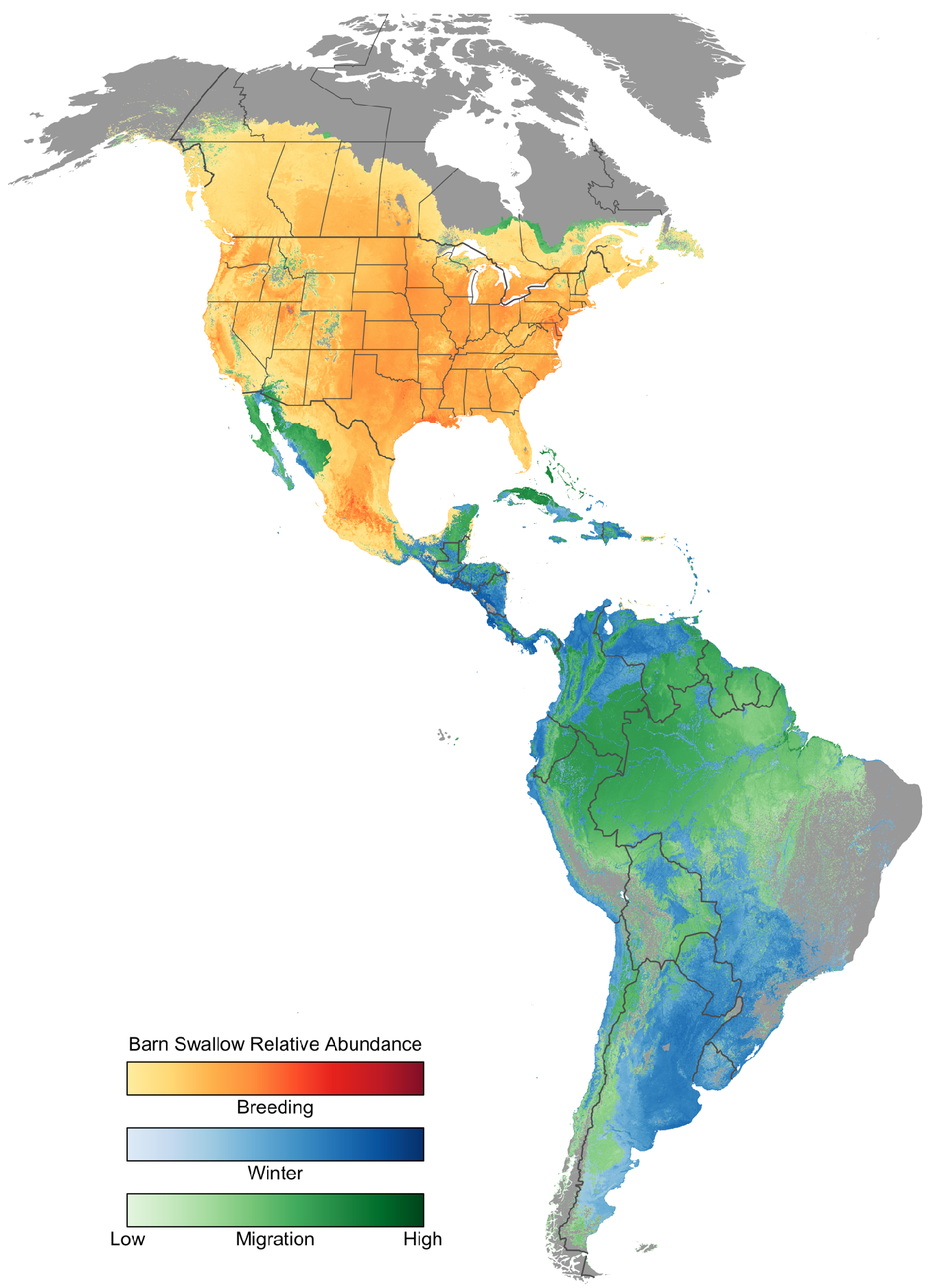
Seasonal Relative Abundance Map for Barn Swallow (*Hirundo rustica*). This map shows the average relative abundance during each of the stationary breeding (June 13-July 20), stationary non-breeding (December 21 - February 8), and non-stationary migratory seasons. The stationary breeding and non-breeding seasons are plotted on top the migratory season, obscuring some aspects of the species’ movements through the annual cycle.

**Figure 7.**
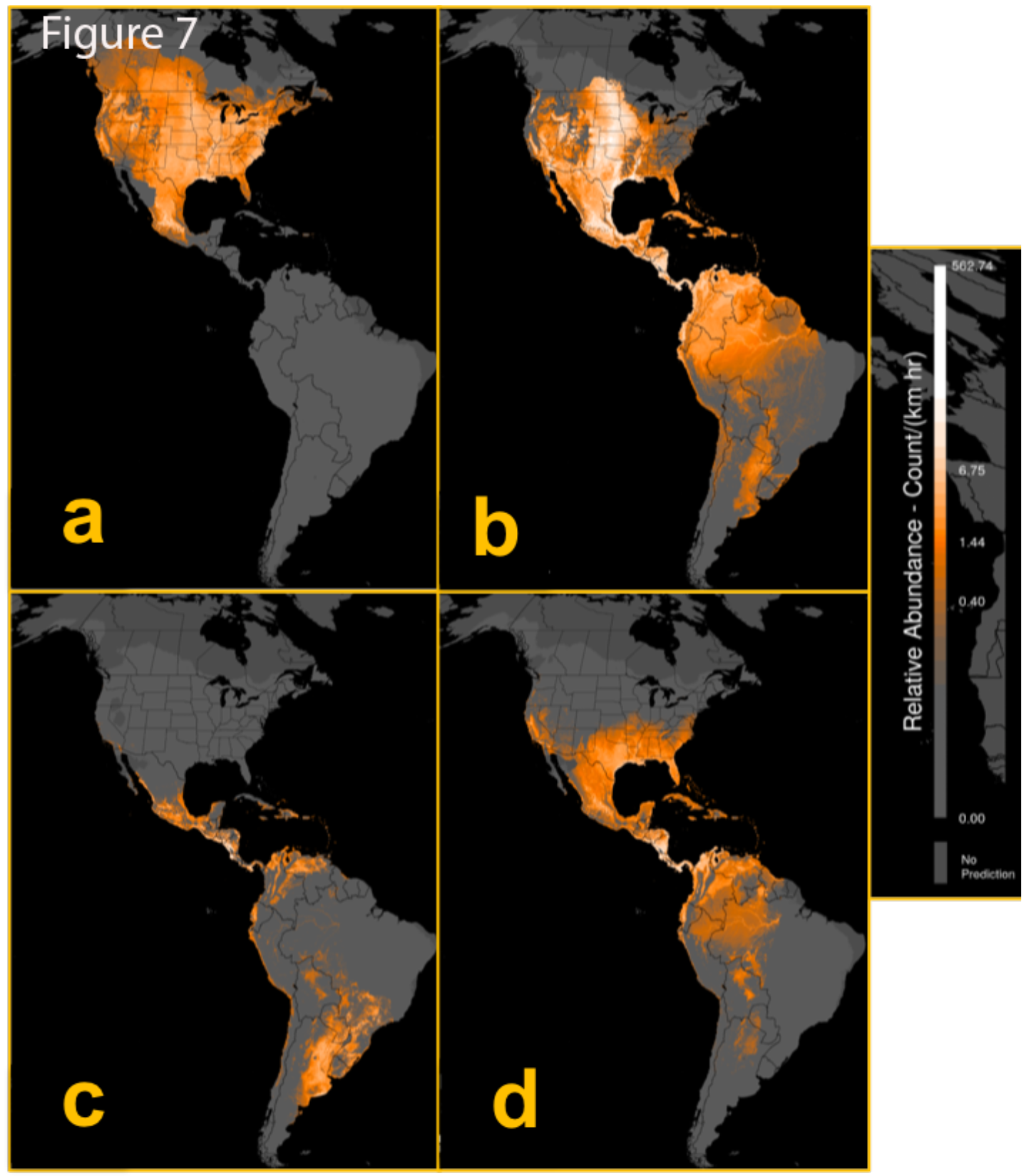
Barn Swallow estimates of relative abundance at 2.8km × 2.8km resolution during (a) breeding (June 21), (b) autumn migration (October 5), (c) non-breeding (January 4), and (d) spring migration (March 28) seasons. Positive abundance is only shown in areas estimated to be occupied with the range boundary depicted as the boundary between pixels with and without color. Lighter orange colors indicate areas occupied with higher abundance. Relative abundance was measured as the expected count of the species on a standardized 1km survey conducted from 7-8AM by a highly experienced participant.

**Figure 8.**
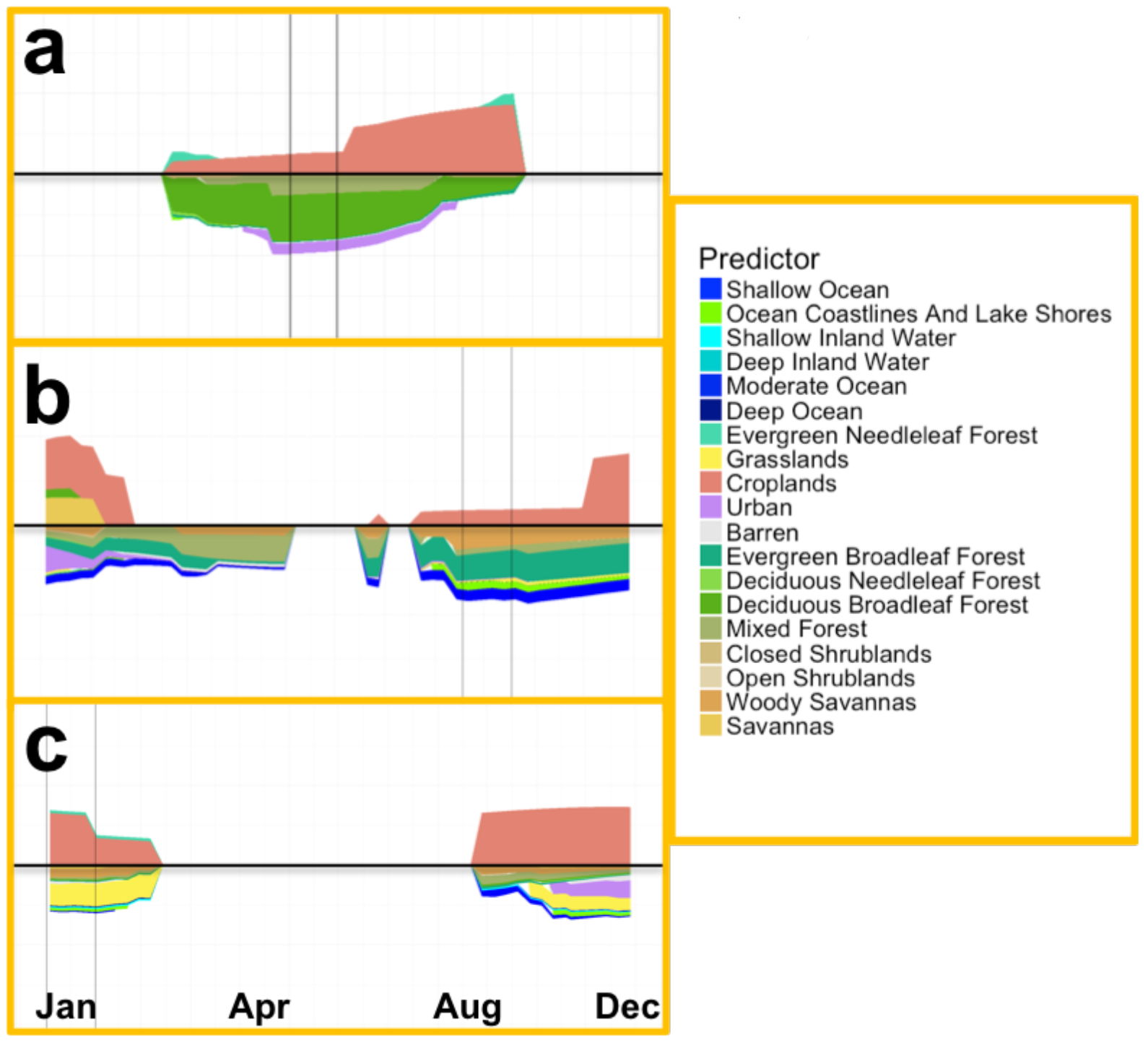
The weekly relative importance of each land and water cover class for Barn Swallow in a) the state of Ohio in U.S., b) Costa Rica, and c) Pampas region of Argentina. Positive importance (colored shading above the horizontal black lines) indicates use of habitat of a landcover type and negative importance indicates class avoidance. The strength of the association with each class is proportional to the height of the class color. Classes with inconsistent association direction, were removed, resulting in total weekly relative importance that sums to less than 1. Barn Swallow shows strong associations with croplands in all regions. In Ohio there is also a positive association with the Deep Inland Water, the class that describes the shore line of Lake Erie, during the spring and autumn migration. In each region, Barn Swallow appears to avoid a different set of landcover classes. This reflects the different landscape structure within which croplands are embedded. In Ohio, croplands are situated within the larger Eastern Deciduous forests with Deciduous Broadleaf Forest, Mixed Forest, and Urban classes. In Costa Rica, there is a mix of Evergreen Broadleaf Forest, Mixed Forest, Woody Savannas, and Oceans, In the Pampas, the landscape is predominantly Grasslands.

Overall, the data contained within eBird complete checklists allow analysts to account for a large proportion of the variation in the observation process. The information on protocol, distance traveled, start time, duration, location, observer expertise, and total number of observers all help us improve our ability to correct for these known sources of variation in detectability. Furthermore, the availability of complete checklists provides a critical source of information on species not detected. Accounting for variation in the observation process gives us greater ability to produce more accurate estimates of the ecological process (Figure 1). This provides greater confidence in trends, maps, and other ecological metrics that are produced using eBird data.

### Ensuring Citizen Science Projects Effectively Monitor Biodiversity

While the fundamental tenets of using citizen science — providing data that meet scientific objectives while ensuring broad participation and improving scientific literacy — are admirable, balancing both objectives can be difficult, and a good balance is not frequently accomplished. We believe that it is possible to engage volunteers in citizen science monitoring via Internet-based means and gather sufficiently robust data to estimate the distributional patterns and trends in species occurrences. While our recommendations include more rigor and standardization in data collection, flexibility in the requirements for individuals to participate in the project can allow for increased participation over time. When the basic components needed to motivate participants are in place, volunteers can be further incentivized to increase participation in ways that are aligned with their motivation. For example, the birding community has responded well to gaming techniques such as leader boards, tools that allow individuals to manage their personal records, and features that allow an individual to explore the patterns of bird occurrences.

Over the next several years, technical advances will improve the potential for an increase in the quantity and quality of citizen science data collected around the world. Already more than 60% of all eBird data are being submitted via mobile apps, which dramatically improve the accuracy of the location and distance an observer walks while observing birds. Developments in the fields of Artificial Intelligence and Augmented Reality, combined with increasingly efficient and powerful mobile technologies, will vastly improve the ability of citizen scientists to detect and identify organisms in the field. For example, projects such as the Cornell Lab of Ornithology’s Merlin Bird ID App, and iNaturalist already use powerful Deep Learning algorithms and computer vision techniques to identify images of thousands of organisms to species, helping observers get to the right species-level identification in the field. Such emerging tools, while powerful, will not be able to fully replace individual expertise, and measured proxies for expertise will remain needed.

Equally important are continuing advances in analytical methodology, which are beginning to provide a means to jointly analyze data collected by both structured and unstructured projects (Fithian et al. 2015, Giraud et al. 2016, Tenan et al. 2016). These models leverage the strengths of both information-rich structured data along with the broad spatial coverage provided by incidental or opportunistic data to improve inferences. These new models can include data collected from different points along the spectrum from unstructured to fully structured.

In summary, the enormous growth of digital networks, rapidly advancing artificial intelligence techniques, the appearance of powerful computing devices that can fit in a pocket, and new statistical analyses will not only improve the quality of citizen science project data, but also improve the inference obtainable from these data. We envision a global network of motivated observers rapidly collecting species lists and observing process information usable for near real-time trend assessment of species and community health, enhanced by real-time information and supported via mobile computing. Whatever the taxon, minimal but critical observation process reporting, as recommended here, will assure improved accuracy in the estimation of the ecological signal. These observations and the process behind them, linked to increasingly improving Earth imagery, promise to support well-designed biodiversity monitoring programs across broad spatial and temporal extents.

